# Evaluating machine learning algorithms to predict lameness in dairy cattle

**DOI:** 10.1101/2024.03.13.584891

**Authors:** Rajesh Neupane, Ashrant Aryal, Angelika Haeussermann, Eberhard Hartung, Pablo Pinedo, Sushil Paudyal

## Abstract

Dairy cattle lameness represents one of the common concerns in intensive and commercial dairy farms. Lameness is characterized by gait-related behavioral changes in cows and multiple approaches are being utilized to associate these changes with lameness conditions including data from accelerometers, and other precision technologies. The objective was to evaluate the use of machine learning algorithms for the identification of lameness conditions in dairy cattle. In this study, 310 multiparous Holstein dairy cows from a herd in Northern Colorado were affixed with a leg-based accelerometer (Icerobotics^®^ Inc, Edinburg, Scotland) to obtain the lying time (min/d), daily steps count (n/d), and daily change (n/d). Subsequently, study cows were monitored for 4 months and cows submitted for claw trimming (CT) were differentiated as receiving corrective claw trimming (CCT) or as being diagnosed with a lameness disorder and consequent therapeutic claw trimming (TCT) by a certified hoof trimmer. Cows not submitted to CT were considered healthy controls. A median filter was applied to smoothen the data by reducing inherent variability. Three different machine learning (ML) models were defined to fit each algorithm which included the conventional features (containing daily lying, daily steps, and daily change derived from the accelerometer), slope features (containing features extracted from each variable in Conventional feature), or all features (3 simple features and 3 slope features). Random forest (RF), Naive Bayes (NB), Logistic Regression (LR), and Time series (ROCKET) were used as ML predictive approaches. For the classification of cows requiring CCT and TCT, ROCKET classifier performed better with accuracy (> 90%), ROC-AUC (> 74%), and F1 score (> 0.61) as compared to other algorithms. Slope features derived in this study increased the efficiency of algorithms as the better-performing models included All features explored. However, further classification of diseases into infectious and non-infectious events was not effective because none of the algorithms presented satisfactory model accuracy parameters. For the classification of observed cow locomotion scores into severely lame and moderately lame conditions, the ROCKET classifier demonstrated satisfactory accuracy (> 0.85), ROC-AUC (> 0.68), and F1 scores (> 0.44). We conclude that ML models using accelerometer data are helpful in the identification of lameness in cows but need further research to increase the granularity and accuracy of classification.

## Introduction

Hoof disorders such as lameness are critical concerns for dairy farms due to economic and animal welfare implications. Lameness not only reduces milk production from cows but also decreases overall cow performance and reproductive capability leading to an increased probability of cows’ removal from the herd [1]. Overall, lameness presents a significant economic loss on farms ranging from $138 and $638 per event, due to costs associated with identification, treatment, and loss in productive performance [2]. Furthermore, lameness is a recognized issue of animal welfare as it represents a behavioral expression of an underlying painful condition [3].

There has been an increased number of cow-based sensors available and used in dairy farms to evaluate animal health and production [4]. As the increasing use of sensor technologies in farms generates large volumes of data, there is a need for the identification of appropriate modern techniques such as Machine Learning to derive useful information from the produced information. In consequence, research efforts using machine learning in the identification and management of animal health and disease are on the rise parallel to the use of sensor technologies. The number of research studies utilizing sensors and machine learning in animal health increased significantly since 2016 in correspondence to the availability of a range of commercial products such as accelerometers for herd monitoring [5].

Early detection of lame cows enables us to timely identify underlying causes [6] and formulate effective treatment strategies. Conventionally, lame cows are identified through visual examination by an expert or herdsman using different indicators in gait assessment such as asymmetric gait, limb abduction and adduction, weight-bearing, arched back, and head bob [7,8]. This detection of lameness through visual inspection holds intrinsic flaws such as intrapersonal and interpersonal biases, efficacy of the observer, and overall subjectivity of the evaluation process. Moreover, farm consolidation and automation have led to a greater number of cattle per employee which demands more accurate and advanced technologies that could assist human cognition for anomalies detection in dairy cattle [9]. Various types of systems including pressure sensors [10], leg-based accelerometers [11], and cameras [6,12] are tested and used for lameness monitoring in dairy farms. However, detection of lameness at a granularity with relevance to management decisions on the farm has not been achieved. Previous studies have used accelerometer-based data for prediction of lameness in dairy cattle and have reported small to moderate association of these behavioral data (lying and standing) for lameness detection [13,14] with an accuracy of 87% [15], sensitivity 90.2%, and specificity 91.7% [16]. Our approach is unique as we aim to predict the cows requiring Corrective Claw Trimming (CCT) or Therapeutic Claw Trimming (TCT) along with lameness scores, which we believe brings more applicable information for on-farm decisions for farmers. Thus, we hypothesized that the evaluation of accelerometer-derived data using machine learning models helps with improved prediction of the lameness events in dairy cattle. Therefore, our objective was to compare various machine learning models for accurate prediction of lameness disease events in dairy cattle.

## Material and methods

### Animal management

The study was conducted at a commercial United States Department of Agriculture (USDA) - certified organic dairy herd in northern Colorado milking approximately 3,500 Holstein cows three times daily, with an average production of 8,600 kg milk per cow during 305 lactation days. The study protocol was approved through the Institutional Animal Care and Use Committee at Colorado State University with IACUC protocol ID 17-7665A. Cows were housed in a free stall barn and had free access to the contiguous dry lot. The average stocking density per number of stalls was 90% for the duration of the study. The barn had a concrete slatted floor with sand bedding and the sand bedding was topped up twice a week. Cows were fed a Total Mixed Ration (TMR) two times a day formulated to meet or exceed the nutritional requirement for a lactating Holstein cow producing 30 kg of milk/d with 3.5% fat and 3.1% true protein [17].

### Lameness management on the farm

Cows on the farm were daily monitored for lameness and cows demonstrating indications of lameness were taken to an examination pen to be assessed for gait-related disorders. All cows on the farm received Corrective (preventive) Claw Trimming (CCT) at least once in six months. The herd under this study had a history of lameness associated with foot rot, white line disease, and digital dermatitis. Those cows that were diagnosed with diseases were treated according to the standard farm protocol by a certified hoof trimmer employed by the farm.

### Data collection

Cows were affixed with an accelerometer on the right rear leg at the fetlock area for the entire duration from 1^st^ December 2017 to 23^rd^ April 2018. After April 23^rd^, cows had access to pasture and to maintain a homogeneous dataset, only records before the grazing period were included in the analysis. The study sample size was limited by the availability of the activity monitoring sensors (a total of 310 sensors). The study included a total of 310 Holstein cows, out of which 69 were primiparous and 241 were multiparous cows, that calved between November 2017 and December 2017. The accelerometer was affixed within 20 days after parturition. The criteria for cow selection were the calving date during the specified period i.e., November 2017 and January 2018, and within 20 days in milk (DIM). Cows were monitored for a week to have adequate system algorithm training before collecting the study data. The activity data (lying time (min/day), steps count (n/day), and daily change (n/day)) were provided based on a proprietary algorithm from the Cowalert system (IceRobotics, Edinburgh, Scotland), summarized in 15-minute intervals, and downloaded onto receivers located in the milking parlor after each milking session. Reference data were collected through weekly monitoring by an experienced claw trimmer, who was positioned at the exit of the milking parlor alley with a full view of cows walking as they returned to their pens, based on different factors such as gait, growth of hoof, cows were classified as CCT or TCT. Furthermore, claw trimmers also monitored the types of lameness on the cows that went for the TCT, briefly illustrated in Table 1. In addition, one of the study personnel (SP) performed biweekly locomotion scoring on a 5-point scale based on Sprecher et al. [18] during the study period. The lameness signs included an abnormal gait, an arched-back posture, or reluctance to put weight on one or more limbs, which would indicate a lameness score of greater than 2 according to Sprecher et al. [18]. These scores were later classified as subclinical cases (scores 2 and 3) and clinical lameness cases (scores 4 and 5), illustrated in Table 1. Locomotion scoring and cows receiving CCT/TCT therapeutic intervention were independent of each other.

**Table 1.** Case definition of different classification terminologies used.

| Case Name | Case definition | Performed by/Classified by | Frequency |
| --- | --- | --- | --- |
| CCT | Routine claw trimming | Claw trimmer | Once every 6 months or need basis |
| TCT | Therapeutic claw trimming and hoof treatment | Claw trimmer | Need basis |
| Moderate lameness | Lameness score (2&3) | SP | Bi-weekly |
| Severe lameness | Lameness score (4&5) | SP | Bi-weekly |
| Infectious lameness | DD, foot rot | Claw trimmer | On the day of CCT |
| Non-infectious lameness | WLD, TLS, Injury, | Claw trimmer | On the day of CCT |

### Statistical analysis and machine learning

Datasets were managed using Microsoft Excel database management software. All the statistical analyses were performed using Python programming language considering various libraries.

### Data preprocessing

The sensor system recorded readings in 15-minute intervals, and daily lying and step data were calculated as a total sum of values from 0000h to 2400h. Records were exported to Excel (Microsoft, 2018) file sheets, and subsequently loaded to the Python programming language (Python Software Foundation, 2023, version 3.10.0). For data reading and imputation, *Pandas* library (version 1.3.0) was used.

To identify and manage outliers, and possible noise in the data from each cow, a 3-day median filter was applied to each of the variables under consideration (daily lying, daily steps, and daily change). Subsequently, feature engineering was performed to derive slope features from existing variables. To calculate slope features, a rolling slope calculation function was created, and the rolling average of the slope was calculated at a window period of 7-day intervals.

A representation of summary statistics of the predictor variables used in the classification of cows into healthy/TCT, healthy/CCT, infectious/noninfectious lameness, and moderately lame/severe lame are illustrated in Fig 1-4. The raincloud plots of the cows comparing TCT vs healthy in Fig 1 indicate higher activity levels in the healthy control cows and a similar observation could be made when healthy control compared to CCT cow (Fig 2). Activity levels for the non-infectious lameness condition were found to be higher when compared to the infectious lameness condition (Fig 3).

**Fig 1.**
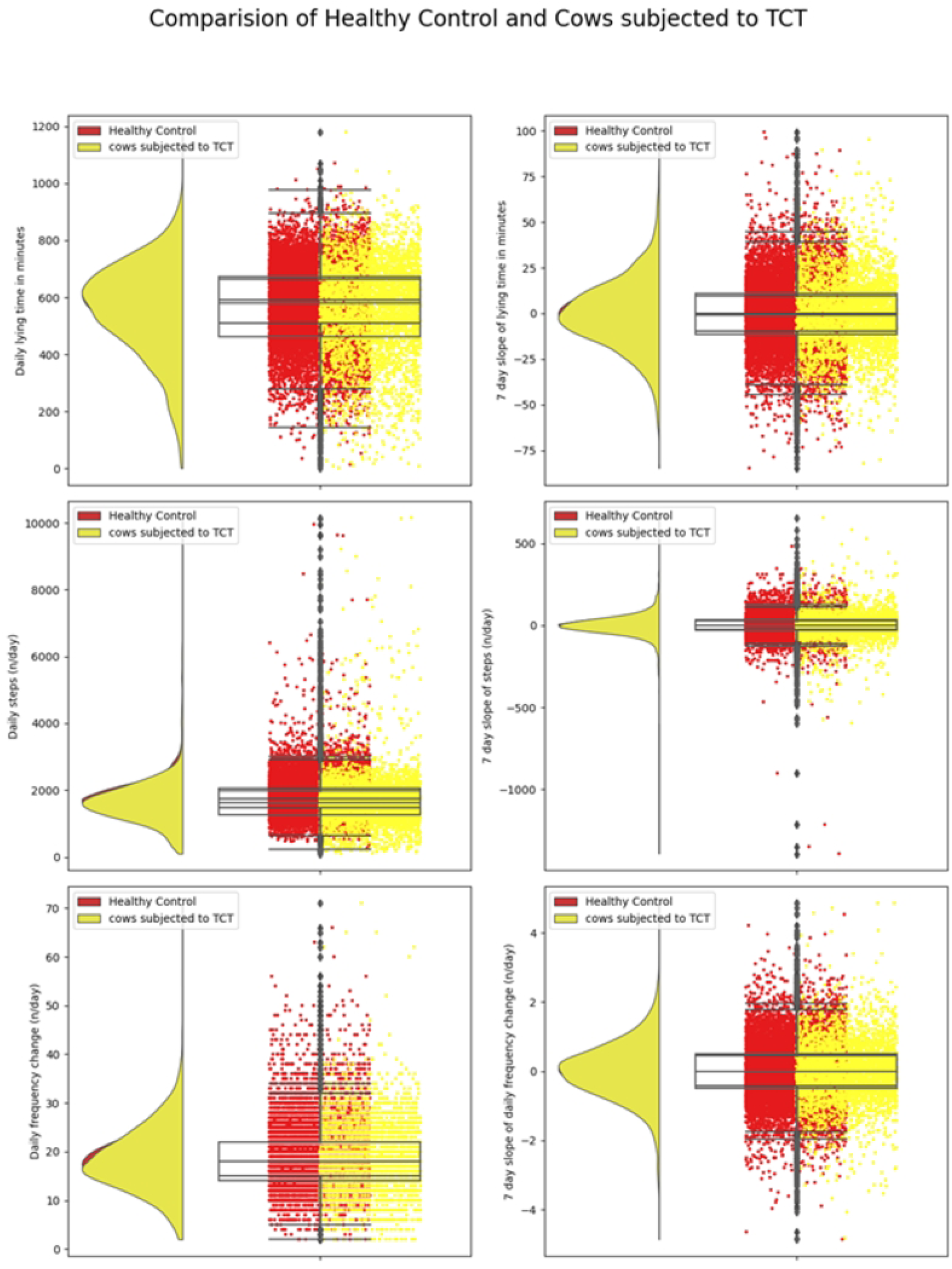
Distribution of daily behaviors and rolling averages of 7-day slopes of these behaviors for cows subjected to TCT and Healthy Control

**Fig 2.**
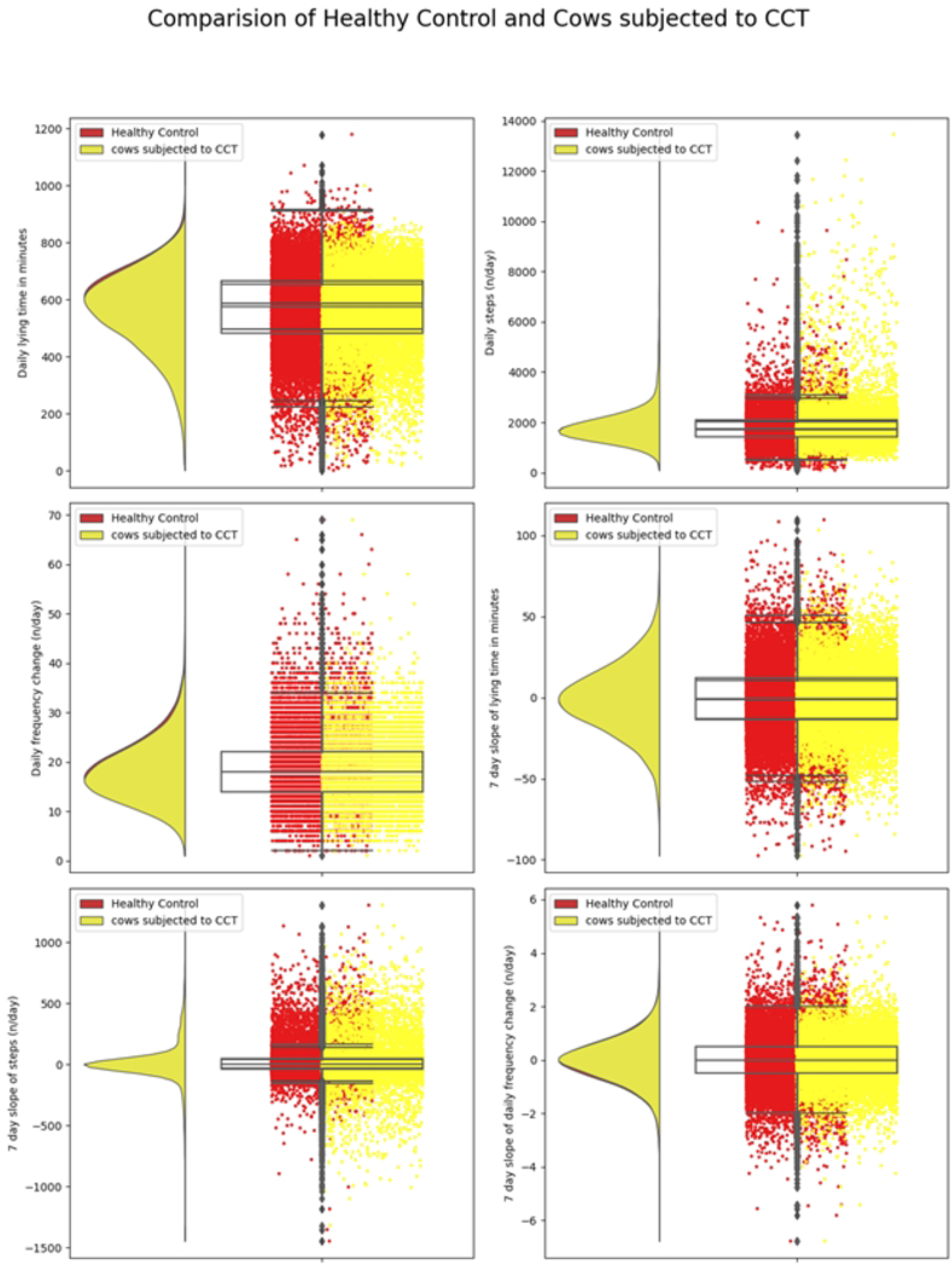
Distribution of daily behaviors and rolling averages of 7-day slopes of these behaviors for cows subjected to CCT and Healthy Control

**Fig 3.**
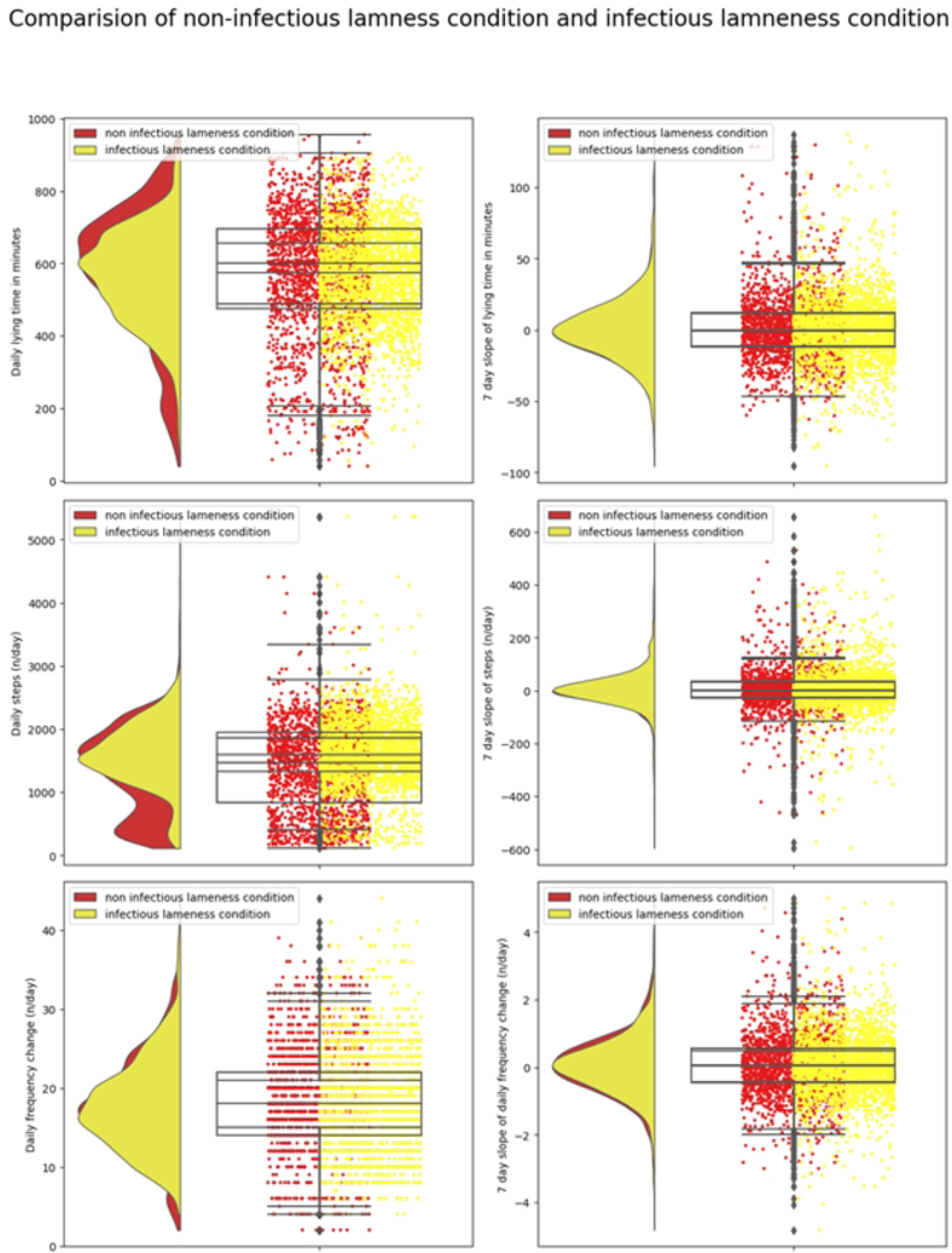
Distribution of daily behaviors and rolling averages of 7-day slopes of these behaviors for cows suffering from infectious lameness and non-infectious lameness.

**Fig 4.**
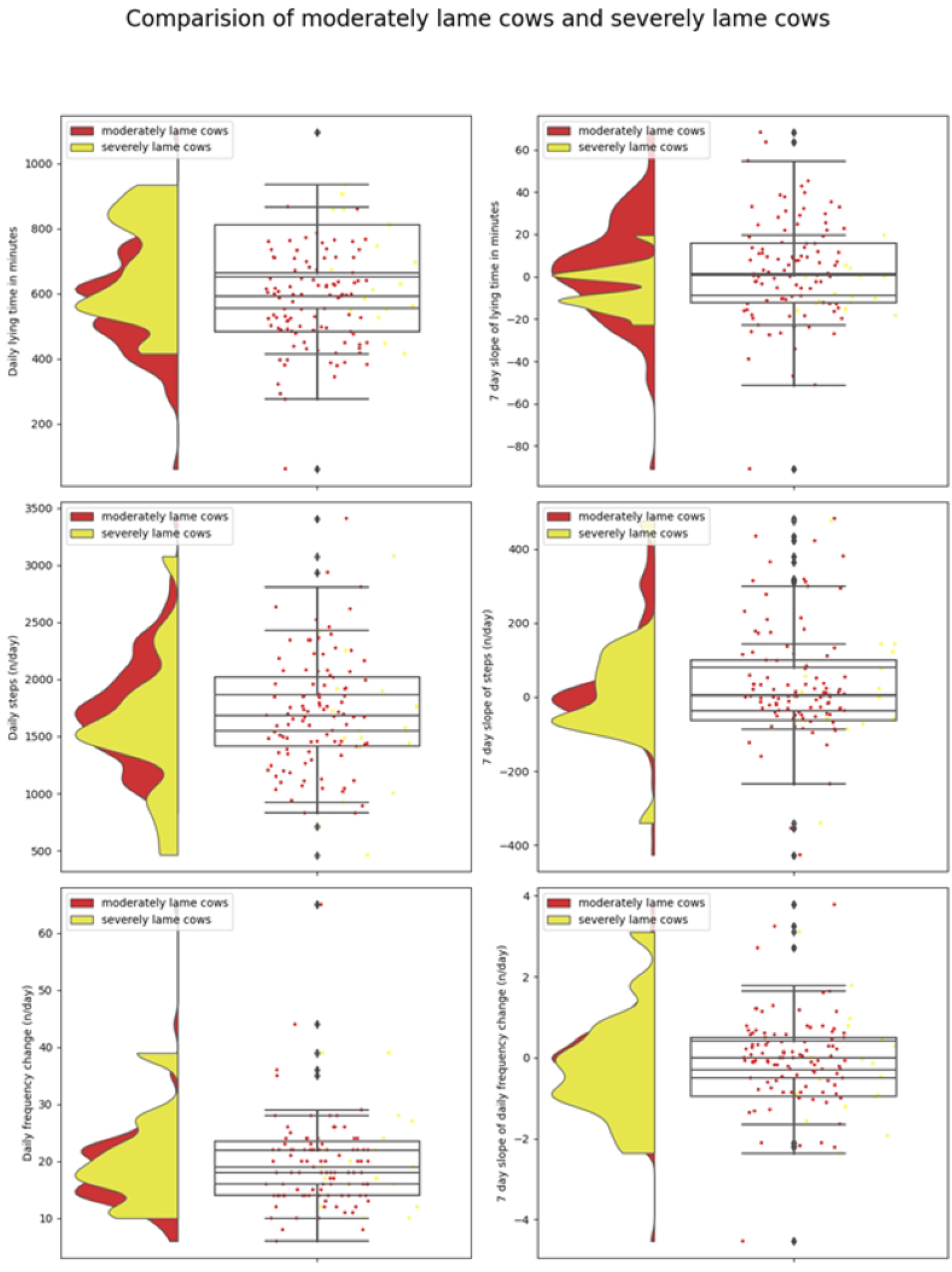
. Distribution of daily behaviors and rolling averages of 7-day slopes of these behaviors for cows affected with moderate lameness and severe lameness condition.

### Definition of outcome variables

Cows were classified as healthy if they did not experience any hoof-related interventions, including corrective or therapeutic claw trimming. Cow events that had confounding effects by generating high activity levels, such as estrus and pen movements, were excluded from the records of healthy cows, specifically within 2 days before and after the detection of an estrus alert. For cows requiring hoof trimming, a dataset was established, encompassing data from 30 days prior to the trimming intervention and 30 days after the event. For the classification of infectious and noninfectious events, daily accelerometer data from each cow suffering from infectious or non-infectious lameness conditions was used. As lameness scores were obtained at every 15-day interval, the daily observations on the scored date were considered for lameness score classification.

### Machine learning model fitting

The Python programming language offers several libraries for machine learning algorithms, which include *TensorFlow, Sklearn, PyTorch*. The latest version of *Sklearn* library (version 1.1) was chosen because of its popularity and easy implementation. The dataset was first loaded in the *Pandas* data frame, and all preprocessing steps were completed. Subsequently, two different approaches were used, one with basic *Sklearn* library where different algorithms such as a Naïve Bayes, Logistic Regression and Random Forest were tested and, with *Sktime library* (Version 0.17.2) ROCKET (Random Convolutions Kernel Transformation) algorithm was tested.

*Sklearn* and *Sktim*e libraries offer the capability of K-fold cross-validation. A 5-fold cross-validation was used in this study, which involves dividing the data into 5 different subsets and training the model on 4 of these subsets while using the remaining subset for testing. This corresponds to an 80% training and 20% testing split. This process is repeated 5 times, using different subsets for testing each time, and the resulting model metrics are averaged, and the standard deviation is reported. Due to the imbalanced nature of the dataset, *Imblearn* (version 0.10.1) library was used to generate samples in the minority class using SMOTE (Synthetic Minority Over-sampling Technique) using default parameters, only for the training data on each cross-validation fold.

For model fitting, we used three different models containing Conventional features, Slope features, and All features to fit in each machine learning algorithm. The Conventional feature model included accelerometer-derived features namely daily lying time, daily steps and daily change, the Slope feature included 7-day rolling slopes of each feature using feature engineering (daily steps, daily lying, and daily change). The All-feature model included both Conventional features and Slope features. The different machine learning algorithms of interest include Logistic Regression, Random Forest, Naïve Bayes, and ROCKET classifiers.

## Results

### General overview of the dataset

The dataset consisted of 36,860 records from 310 cows, with missing values occurring due to technical issues with sensor data downloads on some of the study days. Out of the total records, 7% were missing values, of which, the missing values of continuous variables (daily steps, daily lying time, daily change) were filled with the median of variables because of a high number of outliers (daily lying 2.23%, daily steps 4.8%, and daily change 1.57%) and mode for binary variables such as trim (cows went for trimming interventions), WLD (White Line Disease), DD (Digital Dermatitis), foot rot, injury (foot injury), ulcer (claw ulcer), and heat alert (estrus alert from Cowalert system), as suggested by Peng L and Lei L. [19]. The dataset was again checked for completeness of data and only those cows with complete observations for more than 100 days were used, resulting in a total of 285 cows in the final dataset. In the final dataset, 31 cows were in TCT group; out of which 15 cows with infectious lameness conditions were 15, and 16 with non-infectious lameness conditions, 106 cows in the CCT group, and 12 cows were both subjected to CCT and TCT.

### Results from different machine learning algorithms and models

#### Identification of Corrective Claw Trimming (CCT) and Therapeutic Claw Trimming (TCT)

Four different machine learning algorithms (Naïve Bayes, Logistic Regression, Random Forest, ROCKET) were tested for three different models (Conventional features, Slope features, and All features). A total of twelve runs are performed to predict classification metrics, and only the best-performing models under each classifier are reported. For the models tested for the classification of cows requiring CCT, the models had accuracy in the range of 0.54 to 0.90, F1 score in the range 0.37 to 0.86 and ROC-AUC score in the range 0.54 to 0.89. The ROCKET classifier performed best when using a Conventional feature model with the scores of accuracies (0.90), F1 score (0.86), and ROC-AUC (0.89), as compared to other three different machine learning algorithms (Naive Bayes, Logistic Regression and Random Forest; Fig 5 and Table 2).

**Fig 5.**
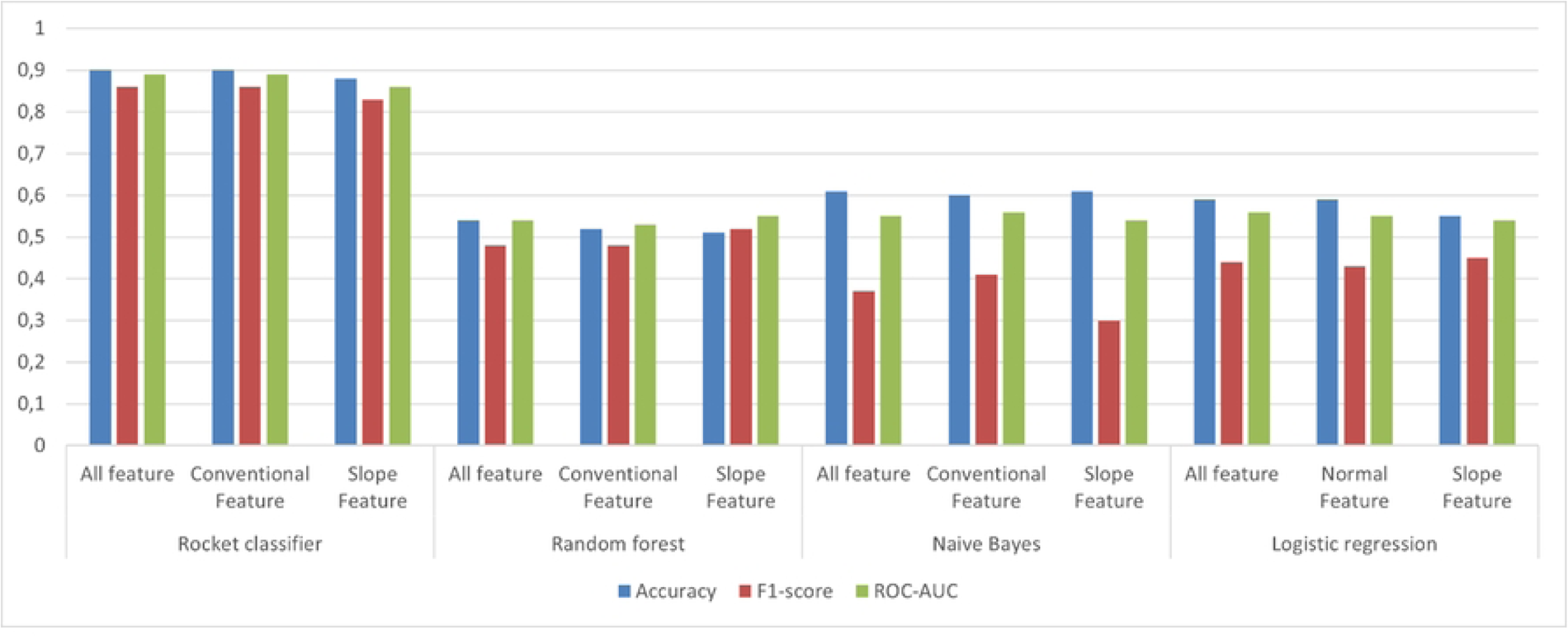
Representation of model performance scores of ML models used for classification of cows needing hoof trimming vs healthy. Conventional feature model consisted of three variables i.e., daily lying time, daily steps and daily change; the Slope feature model included 7 day rolling deviation of each of these variables; All feature models included variables included in Conventional feature and Slope features.

**Table 2.**
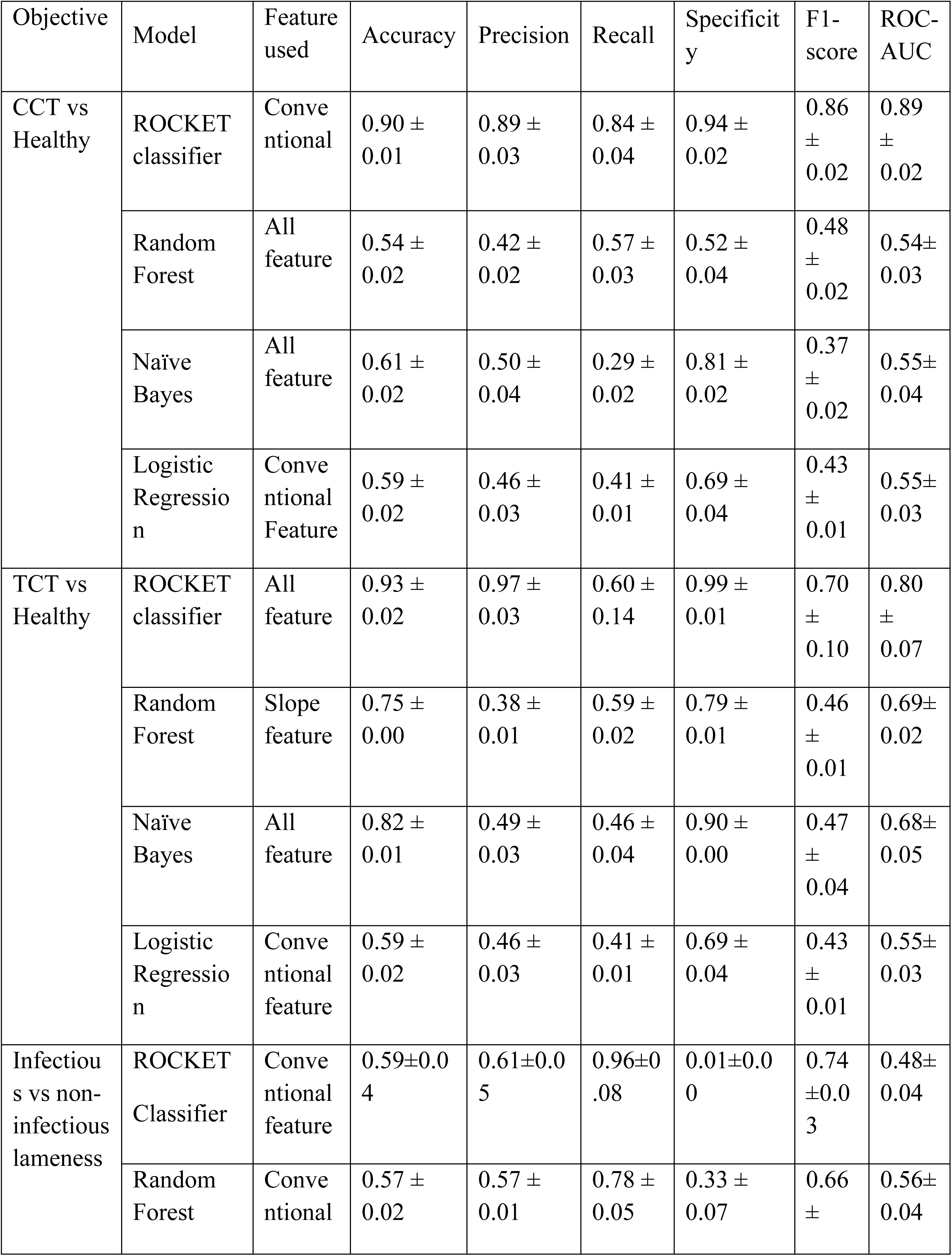
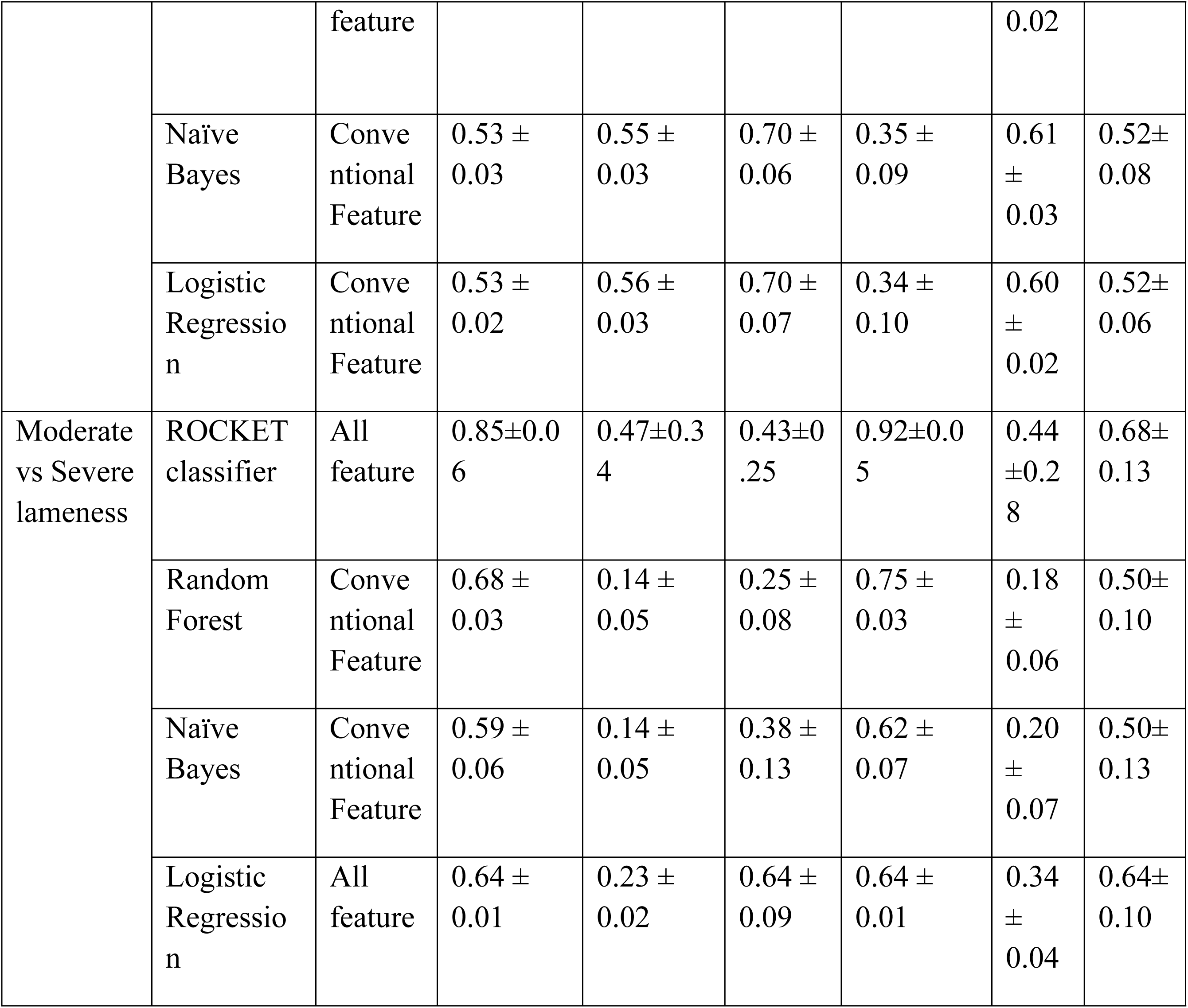
Best performing model scores among the features tested (Conventional feature consisting of daily lying, daily steps and daily change recorded from sensor, Slope feature consisting of a 7-day deviation each feature in Conventional feature, all features consisting of those in Conventional feature and slope feature) for each machine learning models used namely, ROCKET classifier, Random Forest, Naïve bayes, and Logistic Regression.

For prediction of TCT, the models had accuracy in the range of 0.59 to 0.93, F1 score in the range of 0.43 to 0.70 and ROC-AUC score in the range of 0.55 to 0.80. ROCKET classifier performed the best, with all feature model and model scores of Accuracies (0.90), F1-score (0.70), and ROC-AUC (0.80) (Fig 6; Table 2). When compared to the other three different machine learning algorithms (Naive Bayes, Logistic Regression, and Random Forest), none of the algorithms performed significantly well in this evaluation. However, the all-feature model was selected by most of the algorithms as the best-performing combination of features.

**Fig 6.**
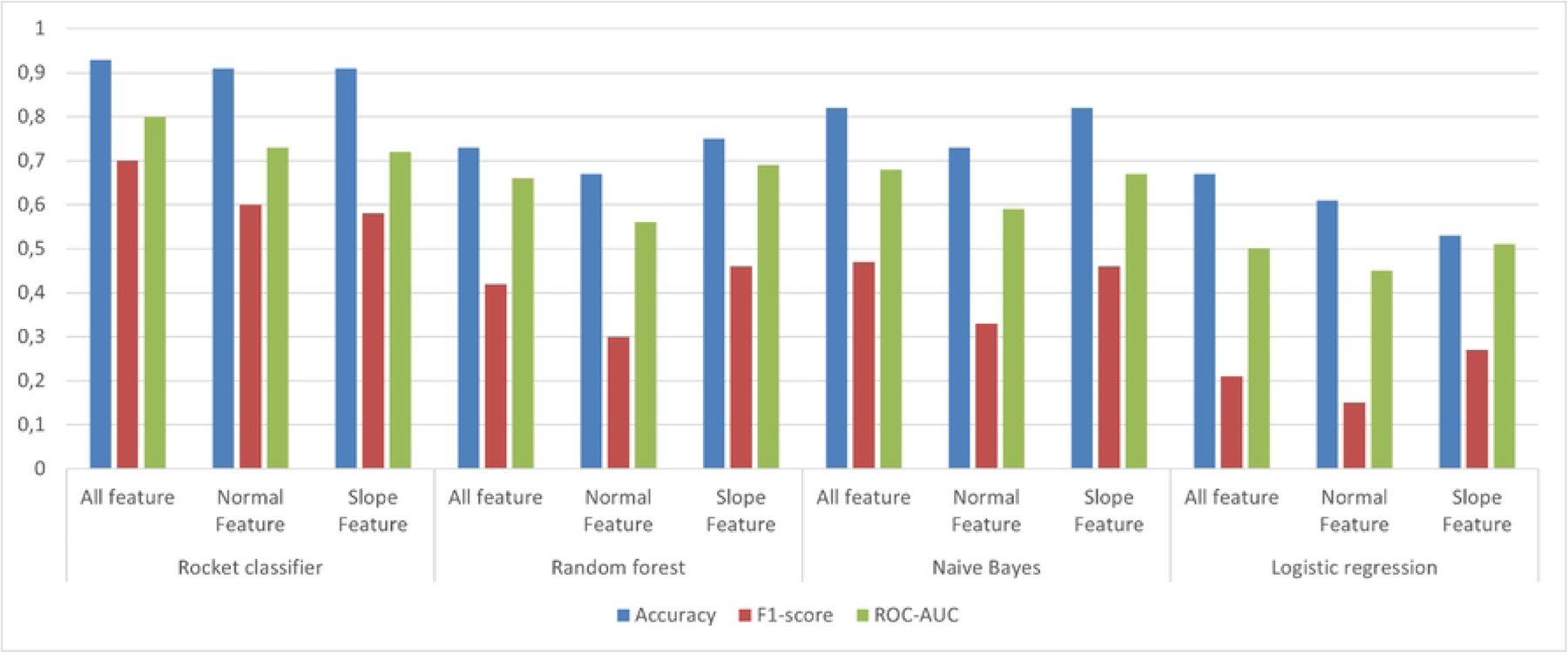
The representation of model performance scores of ML models used for classification of cows needing therapeutic correction vs healthy. Conventional feature model consisted of the three accelerometer derived variables i.e., daily lying, daily steps and daily change; Slope feature model included 7 day rolling deviation of each of these variables; All feature model included variables included in Conventional feature and Slope feature.

#### Classification of diseased cows into infectious and non-infectious lameness conditions

For the comparison of various models for classification of cows into infectious and non-infectious diseases random forest algorithm with Conventional feature demonstrated better performance, with scores of Accuracies (0.57 ± 0.02), Precision (0.57 ± 0.01), Recall (0.78 ± 0.05), Specificity (0.33 ± 0.07), F1-score (0.66 ± 0.02), and ROC-AUC (0.56± 0.04), Fig 7, Table 2. However, Naive Bayes and Logistic Regression performances were similar to Conventional feature with scores of accuracies (0.53), precision (0.55), recall (0.70), and ROC-AUC (0.52) approximately. Furthermore, the performance of the rocket classifier was not satisfactory with low values of sensitivity (00) and ROC-AUC (46) with all feature models.

**Fig 7.**
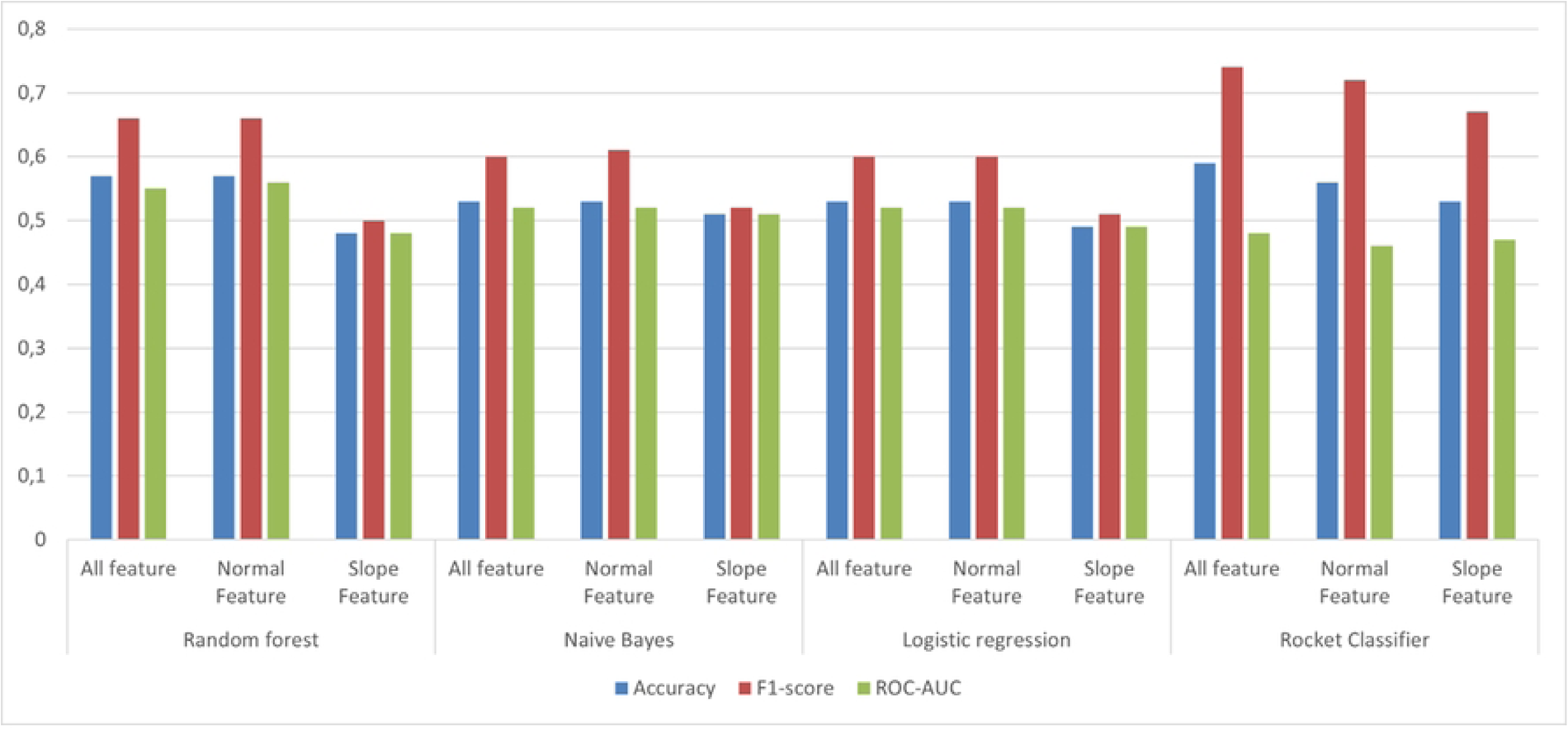
The performance of different algorithms for classification of cows whether they have infectious lameness or non-infectious lameness condition among the diseased cows. Conventional feature model consisted of three accelerometer derived variables i.e., daily lying, daily steps, and daily change; Slope feature model included 7 day rolling deviation of each of these variables; All feature model included variables included in Conventional feature and slope feature.

#### Results for classification of observed lameness scores into moderately lame and severely lame

The model scores for classification of lameness scores into moderately lame and severely lame, were in the range accuracy (0.59-85), F1 (0.18-0.44), ROC-AUC (0.50-0.68), and ROCKET classifier with all features performing better with respect to other algorithms having the scores of accuracies (0.85 ± 0.001), F1-score (0.44 ± 0.28), and ROC-AUC (0.64 ± 0.13) (Fig 8; Table 2).

**Fig 8.**
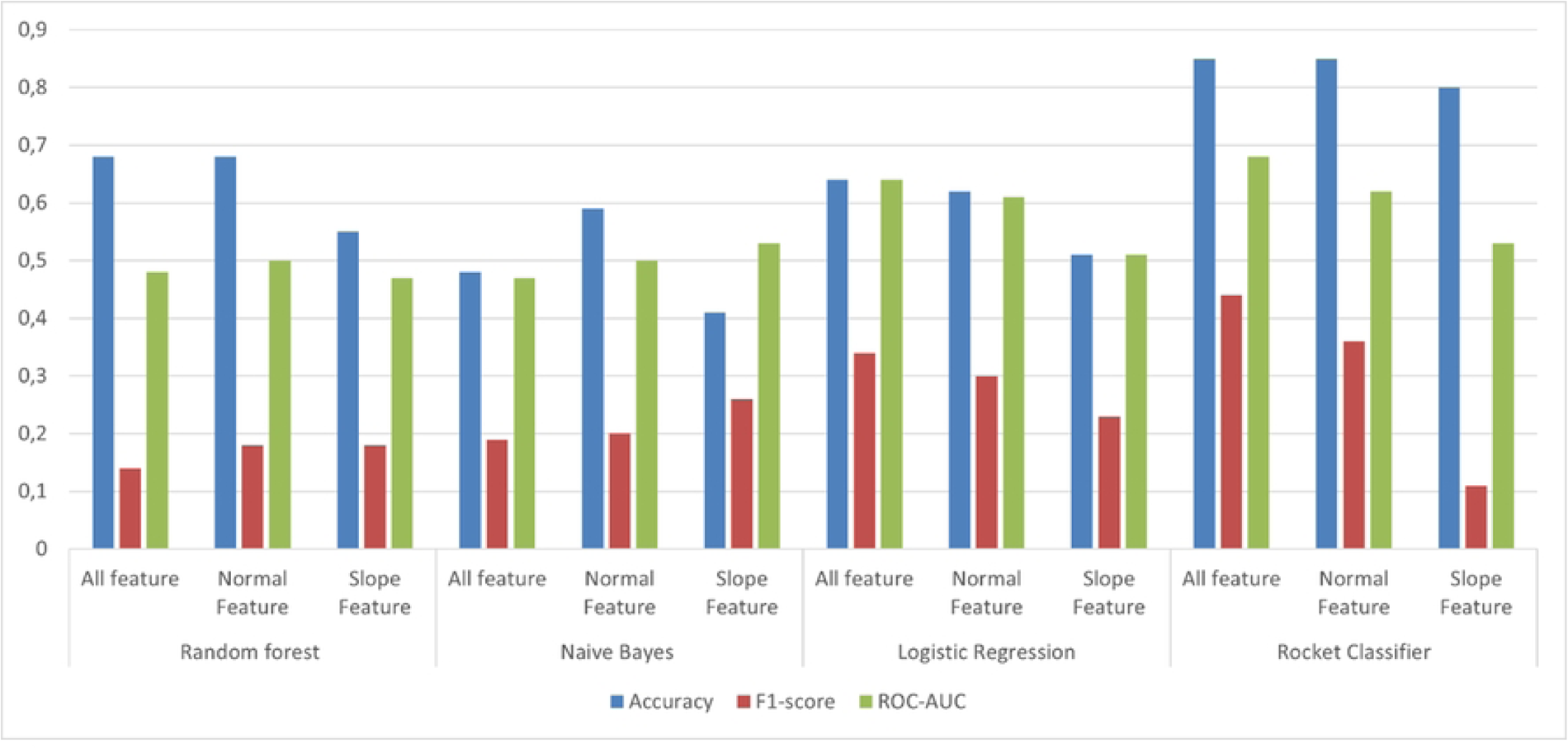
The performance of different algorithms for classification of cows whether they have infectious lameness or non-infectious lameness condition among the diseased cows. Conventional feature model consisted of the three accelerometer derived variables i.e., daily lying, daily steps and daily change; Slope feature model included 7 day rolling deviation of each of these variables; All feature model included variables included in Conventional feature and Slope feature.

## Discussion

In this study, machine learning models were evaluated for the prediction of cows requiring either CCT or TCT based on various accelerometer-derived parameters. The ROCKET classifier demonstrated superior performance for both CCT and TCT classification. Additionally, Random Forest with Conventional features excelled in classifying infectious and noninfectious lameness. Finally, Logistic Regression proved effective in categorizing lameness scores into moderately lame and severely lame conditions.

There are various algorithms available for analysis based on machine learning approach, the popular ones are Logistic Regression, Naive Bayes, Random Forest, and Multilayer perception model. The efficacy of each of the algorithms depends upon the internal working mechanism and nature of data, for example, normalized or unprocessed [20]. In this evaluation, preprocessing of data plays an important role. The different processes such as normalization, removable of outliers, and median filtering have some effects on the results of an algorithm. Working with the missing values in a study to predict lameness, Shahinfar and colleagues [21] suggested that imputing the missing value of rows up to 50% is acceptable and can be imputed from the data respectively. However, another study by Acuna and Rodriguez [22], emphasized the importance of missing values in a dataset and highlighted that the level of missing values is a crucial factor in deciding the appropriate approach for imputation. Their research suggested that datasets with a low percentage of missing values, below 1%, may not have a significant impact on the accuracy of knowledge discovery tasks; datasets with moderate levels of missing values between 1% to 5% may be manageable with simple imputation techniques; and for datasets with higher levels of missing values, between 5% to 15%, more advanced imputation methods such as multiple imputation or regression imputation may be necessary [22]. There are various methods suggested for the imputation of missing data such as KNN (K nearest neighbor), MMI (mean and mode imputation), and RI (Regression Imputation).

Another important consideration when using machine learning algorithms is model building itself. In our study, we had six different features from the accelerometer sensors namely lying time, lying bouts, transition up (frequency of animal getting up), transition down (frequency of animal lying down), and standing time, and animal characteristics such as body weight, days in milk, and other relevant features. In the initial model-building steps, models with all available features were used, but random forest models tended to put more emphasis into characteristics not of specific interest to the study (e.g. DIM and parity). Therefore, variables relevant to the study objectives were chosen from sensor data and were used in model building.

In the pre-processing steps of the sensor dataset, a 3-day median filter was applied to the dataset to remove the large spikes and noise in data as done by Aryal et al., to preprocess sensor signal of heart rate, and skin temperature for fatigue monitoring in construction workers [23]. However, there are other filtering techniques such as low pass filter which discards high frequency and reduces noise from data [24] being used in the study for lameness detection in dairy cattle.

In steps related to feature engineering, a study by Pavlovic and colleagues [25] used *tsfresh* python package to extract features from accelerometer-derived data and subsequently used for behavioral classification. The *tsfresh* package output was used as a guide to develop features used in this study and hence, rolling slopes of 7 days of observation for all features were calculated and the resulting feature was considered a “Slope feature”. We believed that it was a simple yet effective method to capture behavioral differences between lame and non-lame cows. Subsequently, ROCKET classifier was used, which also works internally to produce new features from the data and respectively use these features for classification.

A two-step classification method for the prediction of lameness in sheep was used by Kaler et al., where sensor data were first classified into their respective activities (standing, walking, and lying), and each activity was further classified into lame and non-lame [26]. However, this approach is computationally expensive, and lameness cannot be generalized based on a single activity. Therefore, all three-activity-based variables were used in this analysis. Another concern with this dataset was related to data collected during abnormal activity events like estrus and pen movements. In a recent study, Chapinal et al. removed all the data of cows during estrus period [27]. Accordingly, in our study, 5 days of data (two days before and two days after and day of estrus) of each cow in estrus, were removed.

A study by Byabazaire et al., (2019) uses the clustering of cows into different groups such as active (activity level greater than herd), normal (activity level like herd), and dormant (activity level lower than herd), and derived classification models based on each of these groups. They argued that in bigger herds cows tend to form associations with each other and with this approach, they were able to reduce the classification error by 8%, compared to one-size-fits-all model. Some of our models may resemble a one-size-fits-all model but the ROCKET classifier algorithm utilizes temporal analysis of time-series data within the study period in an individual animal. The evidence from our results also favors that rather than using a one-size-fits-all model, we should look at the possibility of classification of lameness within similar groups or utilize temporal data of individual animals for the analyses [15].

A popular approach for looking for lameness in cows is by categorization of researcher-identified lameness scores into lame and non-lame. A study by Borghart et. Al.,[28] subdivided the lameness scores into sound (score 1) and unsound (scores greater than 1) to identify cows as sound or unsound cows. However, our approach utilized more granular classification by subdividing it into moderately lame (scores 2 & 3) and severe lame (scores 4 & 5). This provides more applicability to dairy farms as this gives an option for farmers to prioritize the management of cows based on available time and resources.

In this study, measurements from an accelerometer that measures both activity and lying behavior were used to predict the cows needing intervention and to further classify lame cows into infectious or non-infectious and moderately lame or severely lame for appropriate and timely management of these diseased animals. Although the herd-level differences between the various variables (daily steps, lying time, and daily change) were not obvious, the median daily steps were low for cows receiving CCT or TCT, with the difference being more evident for TCT cows (134 steps less than healthy cows). Daily lying time only showed a difference of 12 minutes and 16 minutes for TCT and CCT cows, respectively. There was no difference in the median daily change for both groups of cows. The result from this study is concurrent with a recent study by Blackie et al. [29] where they concluded that the total lying time did not differ between lame and non-lame cows. However, other several studies suggested that lame cows had longer lying time and bout duration as compared to non-lame cows [30–32]. Upon further exploration, we identified that lying time was both extremely high and extremely low for the lame cows providing ambivalent information, as cows lie down longer because raising is difficult, on the other hand, cows do not want to rest as some conditions are painful when lying.

For the identification of cows for CCT, the ROCKET classifier had a better model performance as compared to other classifiers. This is due to the internal working principle of the analysis and the nature of the data used in the analysis. A study by Piette et. al. [33] demonstrated that accurate lameness monitoring is only achieved when individualized lameness detection is used and the model achieved scores of sensitivities of 79%, an accuracy of 82%, and a precision of 36.1% [33]. There is a similarity in the approach taken by Piette et. al. [33] and ROCKET classifier we used as they analyze historical data individually for each cow when trying to detect any abnormalities. This was the main reason it achieved impressive scores of accuracies (0.90 ± 0.01), F1 (0.86 ± 0.02), and ROC-AUC (0.89 ± 0.02). While for the other algorithms (Random Forest classifier, Naïve Bayes, and Logistic Regression) data was fed as in individual instances of day labeled either as positive for CCT or negative for CCT rather than individual cows, it is understood that similar observation of variable in cows for positive for CCT and negative for CCT have led low scores as classifiers could not distinguish between two classes.

With reference to the identification of cows requiring TCT, a similar pattern was identified, ROCKET classifier performed better as compared to other algorithms with the scores of accuracies (0.94 ± 0.01), F1 (0.61 ± 0.08), and ROC-AUC (0.74 ± 0.05). An increase in the performance of ROCKET classifier even compared to CCT could be due to stronger behavioral response (lying time, daily steps, and daily change) of lameness towards diseased condition as compared to physiological hoof growth. Other algorithms also performed well in this analysis, as Naïve Bayes with slope feature had accuracy (0.82 ± 0.00), F1 (0.67± 0.0), ROC-AUC (0.47 ± 0.02), maybe because of the low number of instance of cows with TCT as 31 cows were only subjected to TCT, and hence algorithm could effectively learn from data without significant overlap of instances of TCT and healthy cows. Models for the classification of diseased cows into infectious and non-infectious could not effectively identify the events either of infectious or non-infectious diseased cows. The aim was to enable farmers to better manage infectious cows from spreading the disease and was a very novel approach with no relevant literature found to compare the results. Evaluation of a larger dataset containing more events on infectious and non-infectious diseases will help to achieve this goal.

Automatic activity monitoring has been proposed to replace or supplement labor-intensive and time-consuming methods that require visual observation [34]. The sensor devices are effective in measuring cow behavior and useful in herd health monitoring [35]. Detecting the early stages of lameness remains a challenge which corresponds to farmers underestimating the prevalence of lameness leading to delayed treatment of hoof disorders [7,36]. Therefore, there is a need for automated systems that can identify lameness at an early stage without the need for subjective behavioral observations. Thus, automated lameness detection systems and systems to differentiate CCT and TCT bring value in larger herds that have less time for farmers to monitor the animals, and these systems are identified to be more sensitive than traditional observational methods for lameness detection [9].

## Conclusions

This research aimed at comparison of different machine learning algorithms for the prediction of cows requiring claw trimming interventions, classification of cows suffering from either infectious lameness condition or non-infectious lameness condition, and classification of lameness scores. This study concluded that the ROCKET classifier performed better than other tested ML algorithms for classification of cows requiring corrective and therapeutic claw treatment. The model metrics (accuracy, F1, ROC-AUC) for the ROCKET classifier were higher as compared to the performance of other algorithms.

The derived slope feature was identified to be beneficial as well-performing algorithms had All features in the model, and the slope features performed best while predicting TCT with the Naïve Bayes algorithm. This suggests that further feature engineering could make it easier for algorithms to learn from data and hence improve predicting ability. While the classification of the required type of claw trimming performed well, no satisfactory classification between infections and non-infectious or between severe and moderate lameness was reached.

Overall, the study found that the ROCKET classifier is one of the promising tools for predicting cows that require CCT or TCT, while additional information and algorithms are needed to enable more detailed predictions of the type and severeness of the lameness. Further validation, optimization, and real-world deployment would be needed to assess its field accuracy.

## Acknowledgments

The authors would like to thank Ice Robotics Technologies and the dairy farm unit for their help during the research.

